## Supplemental Information for "Cryo-EM structure of the tetra-phosphorylated R-domain in Ycf1 reveals key interactions for transport regulation"

**Author Contributions:** R.S.A.C and T.M.T designed research; R.S.A.C conducted protein expression and purification, Cryo-EM studies, viability assay, ATPase activity assay, mass spectrometry, data analysis and manuscript preparation. M.S.I.R performed protein purification viability assay and ATPase activity assay. N.K.K benchmarked protein purification and supported cryo-EM studies. R.S.A.C, M.S.I.R and T.M.T prepared the manuscript.

**Competing Interest Statement:** The authors declare no competing interest.

**Classification:** Structural Biology, Membrane Proteins

**Keywords:** ABC-transporter, r-domain, phosphorylation, PKC, heavy metal

**Abstract**

Many ATP-binding cassette (ABC) transporters are regulated by phosphorylation on long and disordered loops which present a challenge to visualize with structural methods. We have trapped an activated state of the regulatory domain (R-domain) of Yeast Cadmium Factor 1 (Ycf1) by enzymatically enriching the phosphorylated state. A 3.2 Å cryo-EM structure reveals an R-domain structure with four phosphorylated residues and a position for the entire R-domain. The structure reveals key R-domain interactions including a bridging interaction between NBD1 and NBD2 as well as an interaction with the R-insertion, another regulatory region. We systematically probe these interactions with a linker substitution strategy along the R-domain and find a close match with these interactions and survival under Ycf1-dependent growth conditions. We propose a model where four overlapping phosphorylation sites bridge several regions of Ycf1 to engage in a transport-competent state.

### Supplementary Figures

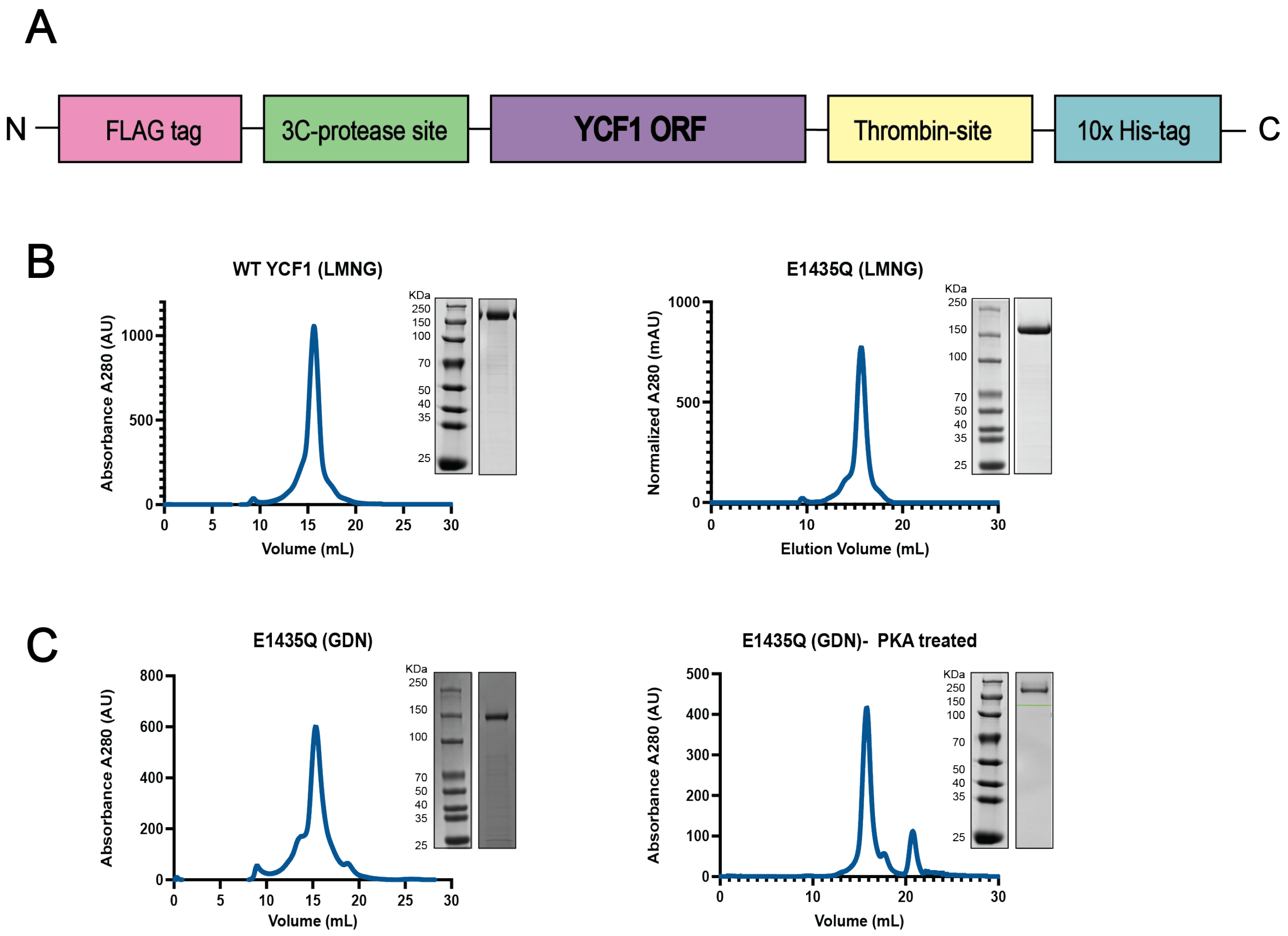


**Supplementary Fig. 1. Ycf1 construct and purification profile.** (A) The DNA construct design for expression of Ycf1 in *S. cerevisiae* utilizing a coupled IMAC affinity with size exclusion chromatography. (B) Size exclusion chromatograms and corresponding 10% SDS-PAGE gel. Wild type and the catalytic dead mutant E1435Q purified in buffer containing 0.01% LMNG/CHS. (C) E1432Q mutant purified in 0.02% GDN detergent and same protein sample after cAMP-PKA treatment to generate hyperphosphorylated state. Hyperphosphorylated samples were then utilized for the preparation of cryo-EM grids and data acquisition.

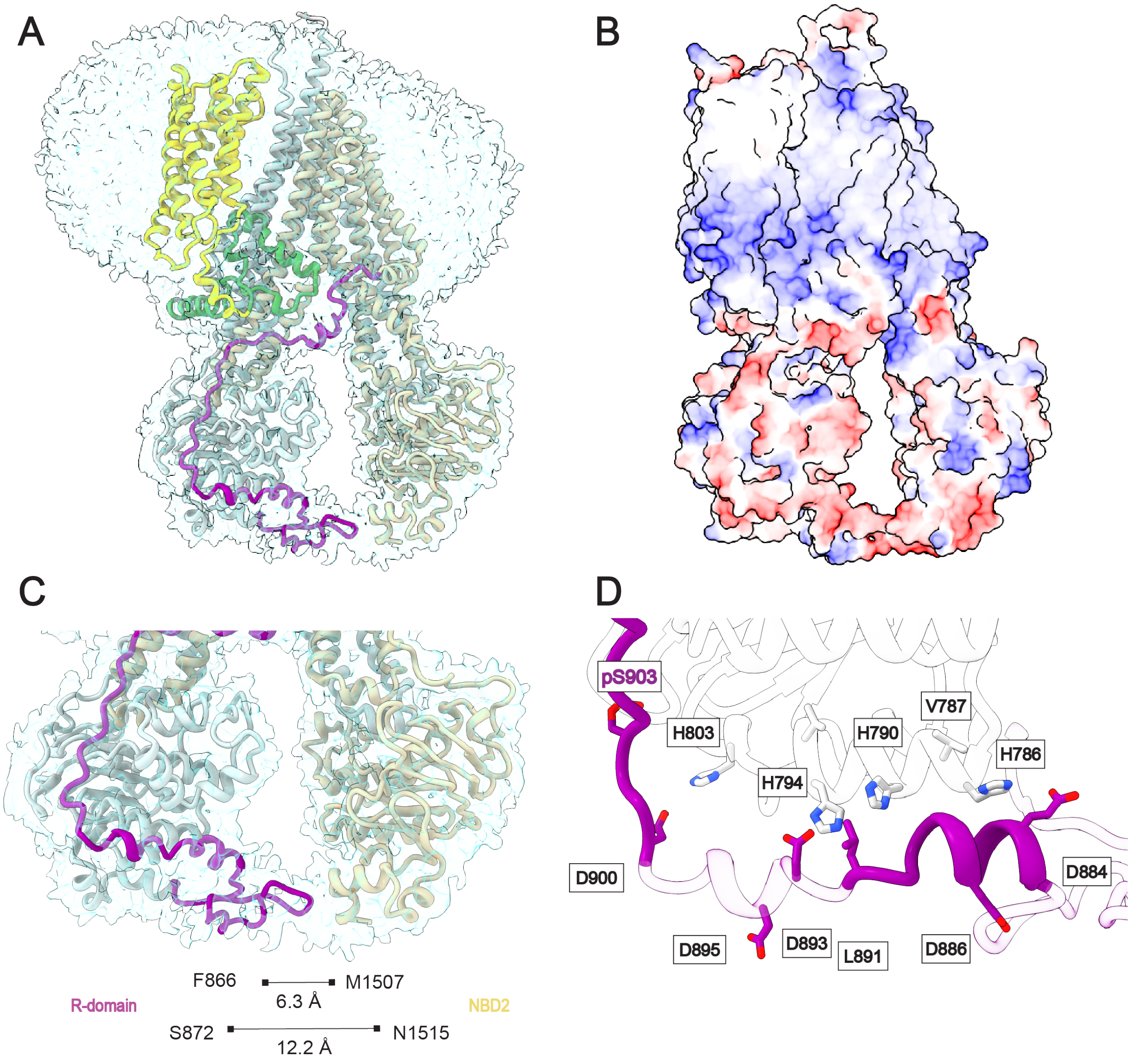


**Supplementary Fig. 2. Local refinement map revealed NBD1 and NBD2 densities in contact.** (A) Overall local refinement map at 3.4 Å obtained from tight masking NBD1. The full R-domain sequence containing continuous low resolution density indicating a potential path towards the C-termini regions. The observation of the R-domain and NBD2 densities in close proximity is a new finding for ABCC family. (B) Electrostatic surface potential (min -24.36, mean 1.23, max 32.50 kcal/(mol·e)) of the local refined map presenting a R-domain segment that is highly negatively charged, even on the lower NBD1-NBD2 portion remains the same overall charge. (C) The R-domain-NBD2 density with hypothetical model of Ycf1 with gaps filled in from the Ycf1 Alphafold structure. (D) Potential electrostatic contacts between the model R-domain position and NBD1. Dark purple represents modeled regions of the R-domain, light transparent purple represents regions left out of the final model but present in the EM density. NBD1 is colored white.

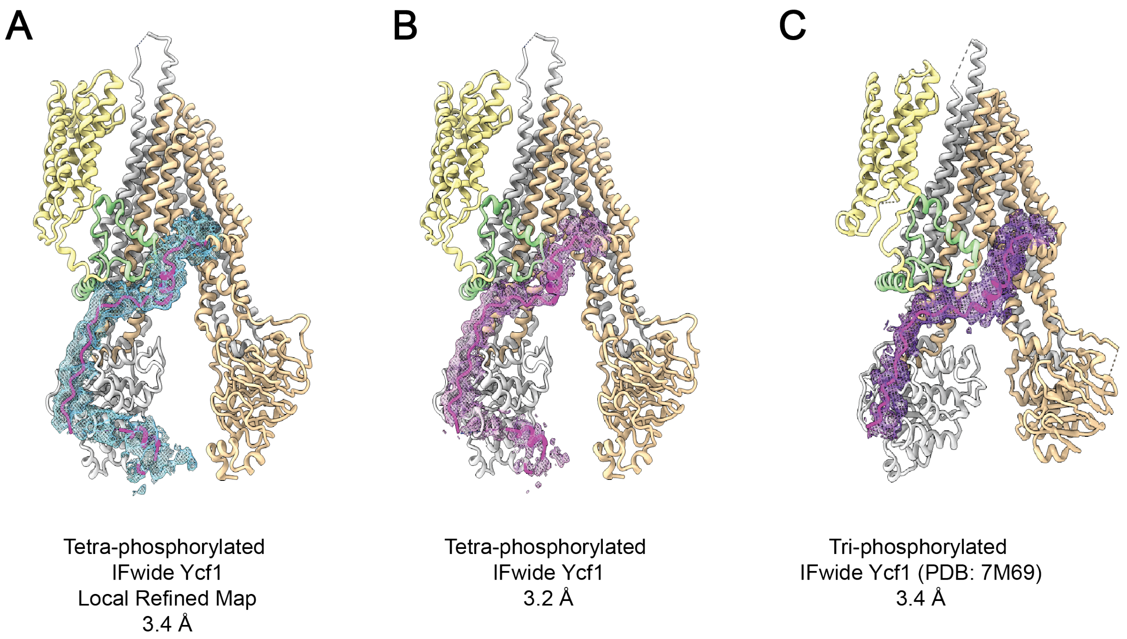


**Supplementary Fig. 3. Improvements in the R-domain coverage on the inward-facing wide Ycf1 structures through cryo-EM.** (A) The local refinement expanded the continuous electron density map from the conventional non-uniform refinement map in (B). Although this region is at the lower resolution range it suggests that this disordered strand can report to NBD2 although in close association with the NBD1. (B) The non-uniform refinement map improved the overall resolution and expanded the R-domain placement. The phosphorylation at serine 903 was only possible due to the expansion of the coverage in the R-domain map. (C) Previously reported wild-type Ycf1 structure (1,2) could not determine the N-termini region of the R-domain due to limited resolution. Maps are presented at step 1 and contour level at 0.014, with the selected R-domain residues 855-935 shown at 2.5 radius surface zone for both local and regular tetra phosphorylated map. The 7M69 map (1) and model were presented similarly with a 0.2 contour level and 2.3 radius surface zone

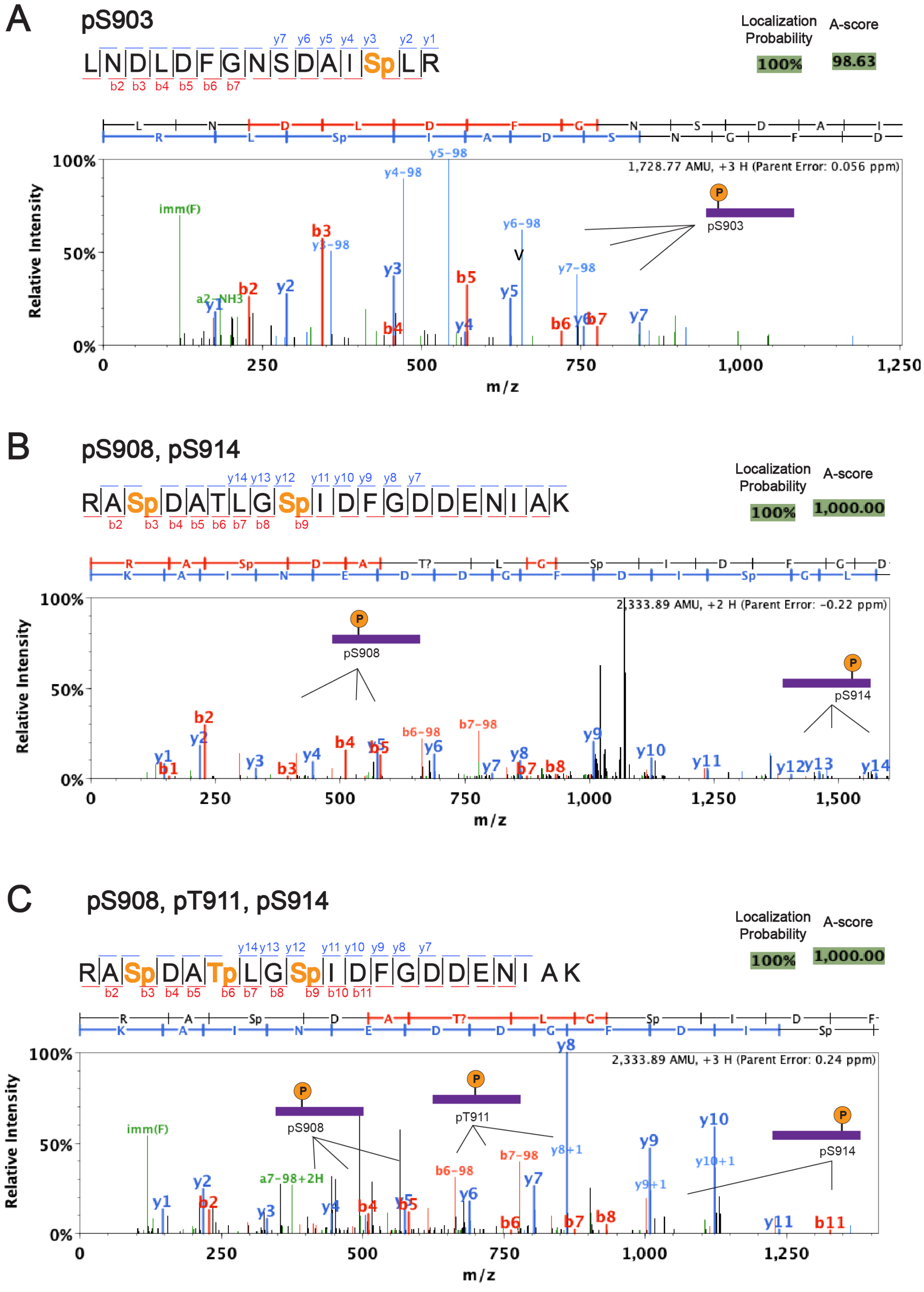


**Supplemental Figure 4. Mass spectrum of the PKA treated Ycf1 confirming the tetra-phosphorylated state in the main sites: S903, S908, T911 and S914.** The identified y and b ions from the PKA phosphorylated Ycf1 are shown below the peptide sequence annotation, in which y ions are presented in blue and b ion in red. Phosphorylated residues are colored orange in the sequence annotation and indicated in the mass spectrum. Localization probability and A-score for the specific peptide sequence is show on the top right corner, similar patterns were found to those reported in the previously published Ycf1 sample (1) .(A) The novel S903 phosphorylation presented multiple confirmatory peaks and it is identified in the peptide ion sequences (B) Additionally the double phosphorylated (pS908, pS914) peptide can be found simultaneously as well as the tri phosphorylated (pS908, pT911, S914. (C) The triple phosphorylated peptide species is often a prevalent population in the Ycf1 mass spectrum studies. Mass spectrometry for phosphorylation identification were run in duplicates.

| Residues | Coevolving Motifs | Probability | Score | Distance (Å) |
|---|---|---|---|---|
| S872 - L912 | PKC-CKII | 0.74 | 1.05 | 57.9 |
| S872 - D909 | PKA-CKII | 0.71 | 0.91 | 52.6 |
| V874 - D909 | PKC-CKII | 0.85 | 1.75 | 51.6 |
| V874 - A910 | PKC-CKII | 0.79 | 1.30 | 52.1 |
| R875 - A907 | PKC-PKA | 0.87 | 1.88 | 44.8 |
| R875 - S908 | PKC-PKA | 0.74 | 1.06 | 49.7 |
| R906 - A907 | PKA-PKA | 1.00 | 8.48 | 5.5 |
| R906 - D909 | PKA-CKII | 1.00 | 6.00 | 9.0 |
| R906 - S908 | PKA-PKA | 1.00 | 5.40 | 7.9 |
| R906 - A910 | PKA-CKII | 0.95 | 3.03 | 13.2 |
| R906 - T911 | PKA-CKII | 0.88 | 1.98 | 16.1 |
| R906 - L912 | PKA-CKII | 0.80 | 1.39 | 17.7 |
| S914 - D920 | | 0.75 | 1.12 | 17.6 |
| E921 - H928 | | 0.82 | 1.50 | 15.2 |
| E927 - V934 | | 0.89 | 2.11 | 17.8 |

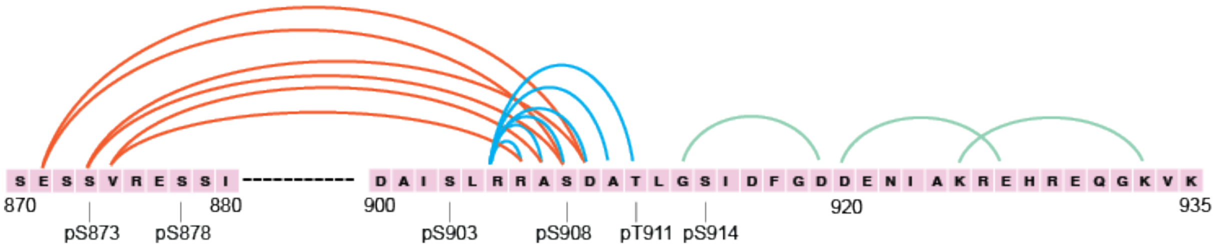


**Supplemental Fig.5 Evolutionary coupling analysis of co-evolving residues of Ycf1 R-domain.** The hydrophobic and charged residues featuring the R-domain N-terminus have coevolved with the core phosphorylation sites as well as C-terminal residues as was also shown in (1). Strikingly, the kinase motifs for the phosphorylated residues present strong co-evolution to the key regulatory sites.

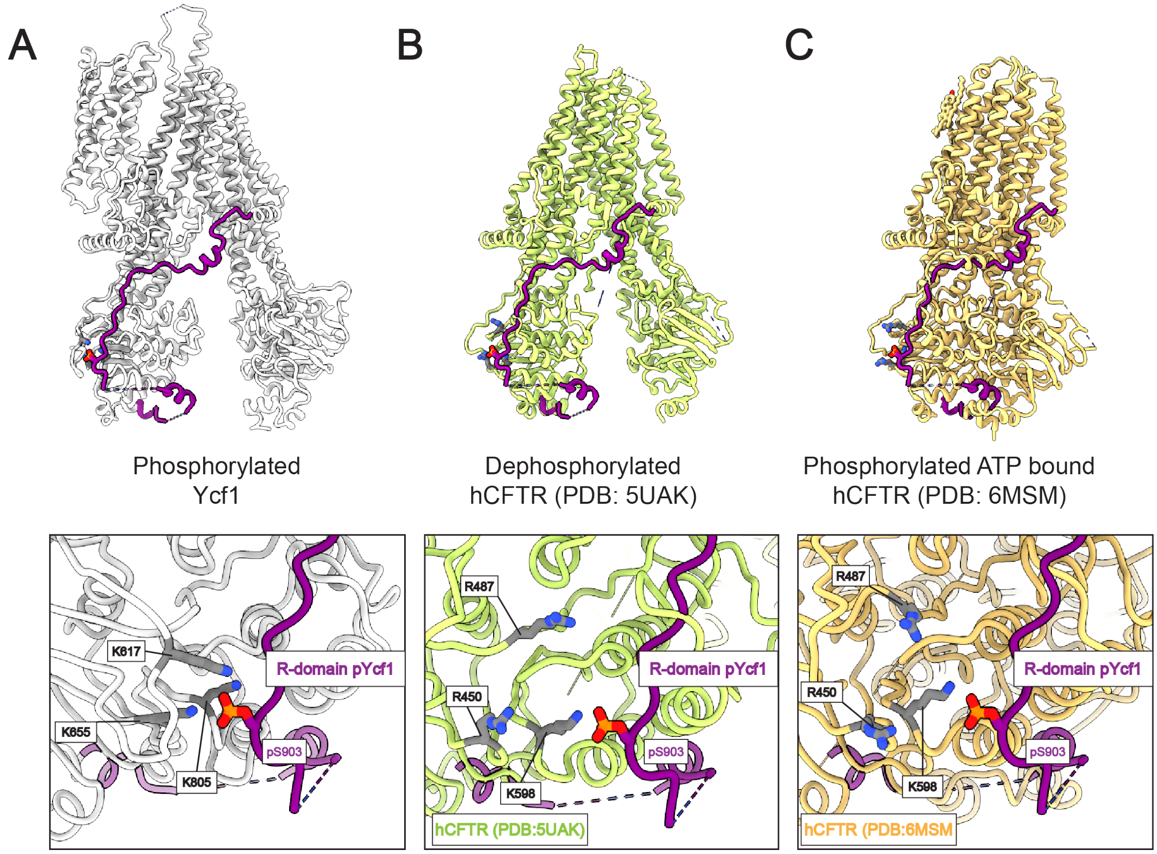


**Supplemental Fig. 6 Ycf1 and CFTR share similar binding pocket for phosphorylated S903.** The S903 site from the tetra phosphorylated Ycf1 in its binding site presented in this study is show in silver (A). Structural comparisons of current CFTR structures with the superimposed R-domain Ycf1 from this study presents a common binding interface. (B) The dephosphorylated CFTR (3) still resembles the pocket with similar chemistry, with the replace of lysines (Ycf1 K617 and K655) for two arginine residues (CFTR R450 and R487). (C) The ATP bound and phosphorylated CFTR structure (4) although shown in a dimerized state is still able to retain its binding pocket. Although these basic sites are not conserved in multiple sequence alignments among ABC-C transporters, this structural comparison provide insights in this phosphorylation interacting hotspot.

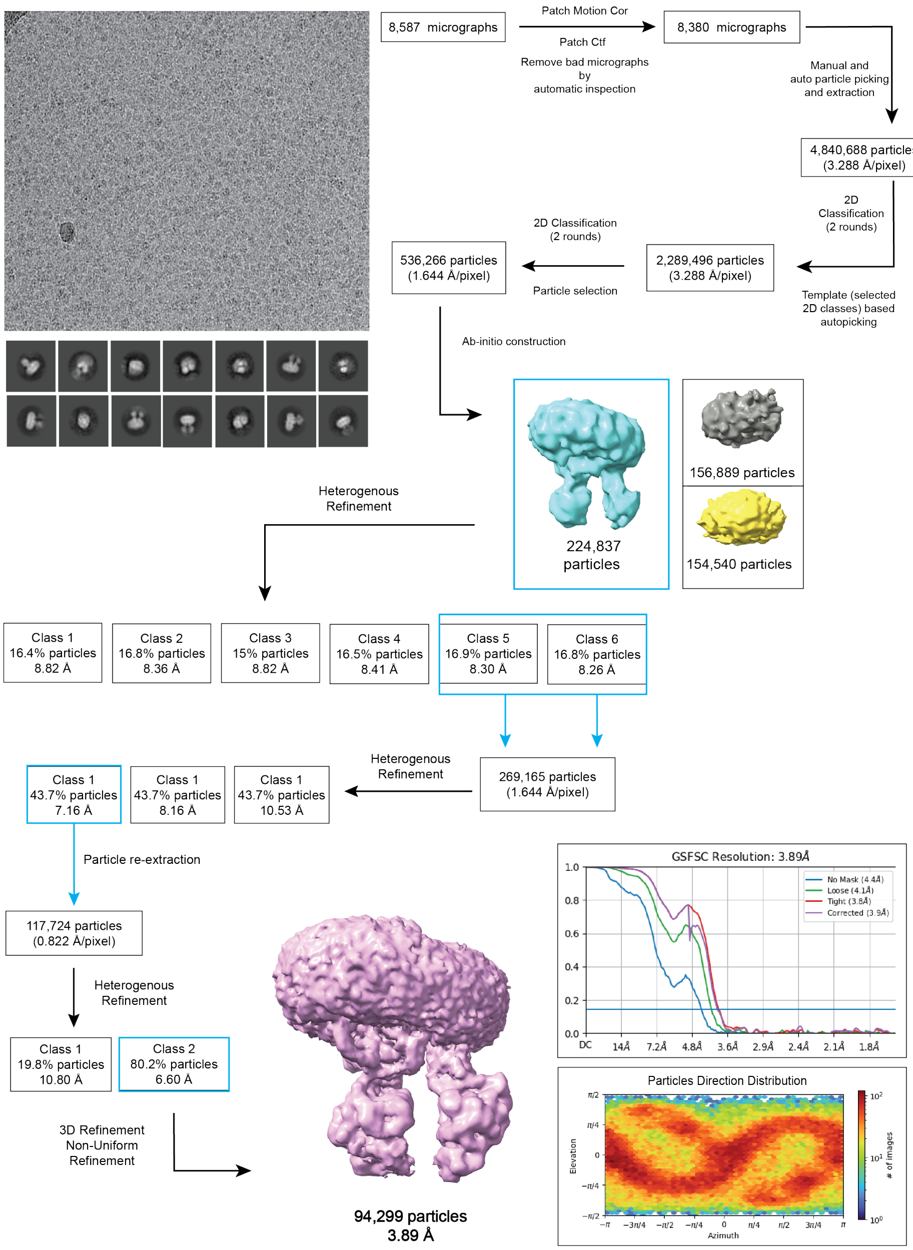


**Supplemental Figure 7. Cryo-EM data processing workflow for E1435Q PKA-treated Ycf1.** The electron microscopy image processing workflow for the initial dataset utilizing CRYOSPARC. This initial first refined map at 3.89 Å was used as 3D template for deep learning particle picking and another round of processing.

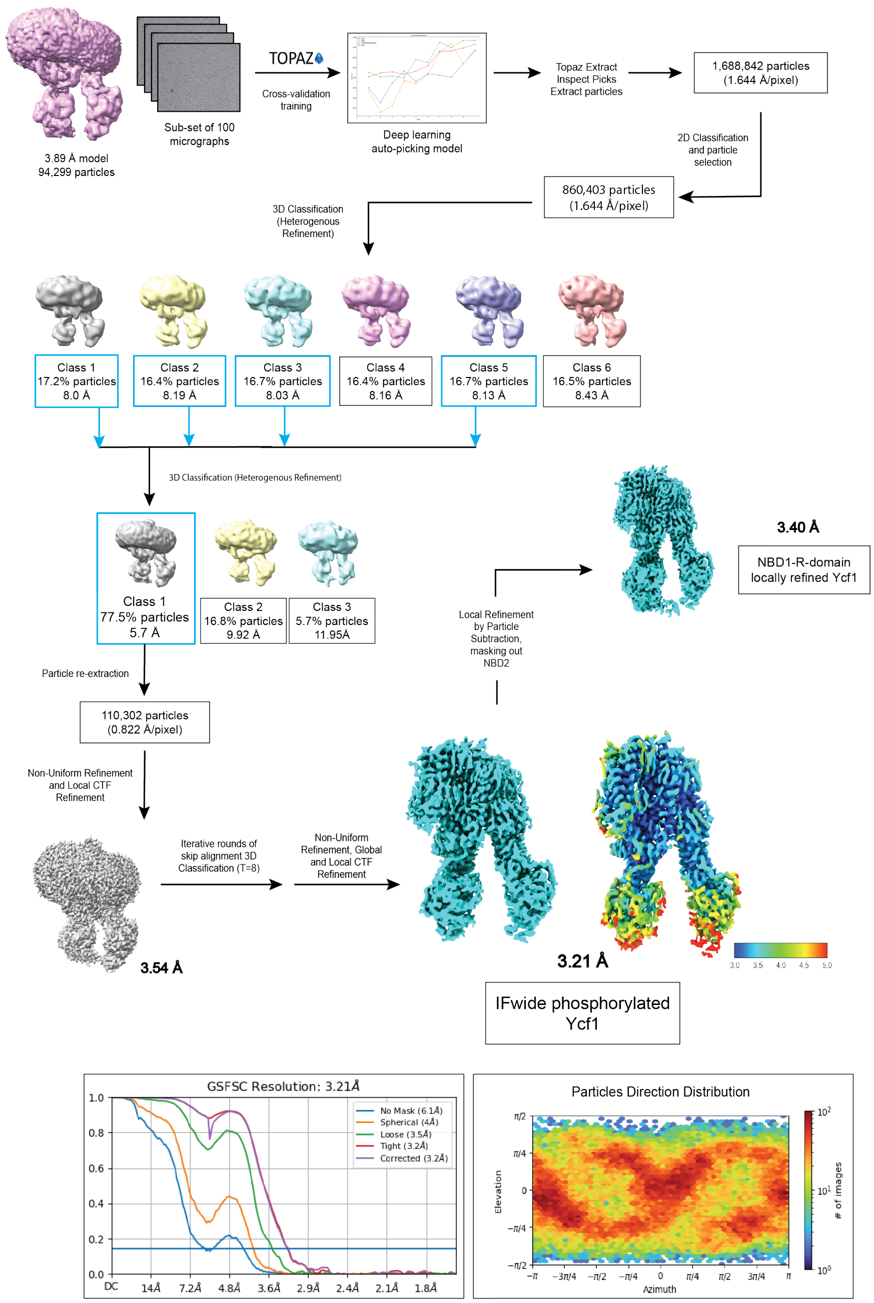


**Supplemental Figure 8. Cryo-EM reference based data-processing workflow for E1435Q phosphorylated Ycf1.** Pipeline utilized in image processing using a previously generated 3D model to improve final map resolution. Artificial intelligence for assisted particle picking (5) in an interactive way in order to optimize auto-picking models and consequently generate improve accuracy in particle detection. Further, the optimized particle set is re-processed and indeed it was possible to obtain a significant improvement in the refined map resolution as well as global features and side-chain sharpening.
